# Grazers and predators mediate the post-settlement bottleneck in Caribbean octocoral forests

**DOI:** 10.1101/2022.08.09.503401

**Authors:** Christopher D. Wells, Joseph Benz, Kaitlyn J. Tonra, Emily R. Anderson, Howard R. Lasker

**Affiliations:** Department of Geology, University at Buffalo, State University at New York, Buffalo, NY, USA; Department of Environment and Sustainability, University at Buffalo, State University at New York, Buffalo, NY, USA; Department of Integrative Biology, Oregon State University, Corvallis, OR, USA; Smithsonian Environmental Research Center, Edgewater, MD, USA

## Abstract

Caribbean octocorals have not suffered the decades long decline in abundance that has plagued reef-building scleractinian corals. Their success and the formation of octocoral forests has been attributed to their continuing recruitment to reef habitats. Assessing the processes controlling recruitment is essential to understanding the success of octocorals and predicting their future. Benthic grazers on coral reefs can facilitate the growth and recruitment of corals by reducing the abundance of competitive algal turfs and macroalgae or hinder corals through predation of coral tissue and recruits. We assessed the effects of grazing by fishes and the sea urchin *Diadema antillarum* and mesofaunal predation on octocoral recruitment in a series of manipulative experiments using varying grazer/predator exclusion and inclusion conditions in *in situ* and *ex situ* experiments. Exposure to fish and urchin grazing significantly reduced survival and recruitment of single-polyp octocorals, while turf-associated mesofauna did not significantly affect neither recruitment nor survival. We also found a positive relationship between octocoral recruitment and turf algae, a potential related response to the deleterious effect of grazing exposure. These data suggest that grazers and predators mediate the mortality bottleneck characteristic of recruitment. Thus, the declines in the abundance of grazing fishes and urchins throughout the Caribbean may have contributed to the increase in abundance of octocorals in the Caribbean, concurrent with the loss of scleractinians.

## Introduction

As scleractinian corals have declined globally (Gardner et al. 2003; Bellwood et al. 2004; Hughes et al. 2018), octocorals have emerged as one of their major successors on Caribbean reefs (Norström et al. 2009; Lenz et al. 2015; Edmunds and Lasker 2016; Sánchez et al. 2019). Octocoral forests provide some of the same functions as scleractinian reefs (e.g., Privitera-Johnson, Lenz, and Edmunds 2015; Tsounis, Steele, and Edmunds 2020) and are resilient to climatic events like hurricanes (Lasker et al. 2020) unlike Caribbean scleractinians (Gardner et al. 2003). Octocoral success has been attributed to reduced direct competition with scleractinians, higher fecundity, and greater juvenile survivorship and recruitment (Ruzicka et al. 2013; Lenz et al. 2015; Bartlett et al. 2018; Lasker et al. 2020; Tsounis and Edmunds 2017; Martínez-Quintana and Lasker 2021). While these mechanisms are key to understanding shifting reef communities, the finer details of octocoral recruitment processes remain understudied compared to those of scleractinians.

Macroalgae have often been highlighted as a threat to juvenile recruitment of both scleractinians and octocorals (Wells et al. 2021; Linares, Cebrian, and Coma 2012; Kuffner et al. 2006; Hughes 1994; Bruno et al. 2009; Roff and Mumby 2012). Proposed mechanisms for this negative interaction include allelopathic competition (Kuffner et al. 2006; Rasher et al. 2011; Bonaldo and Hay 2014), algae providing a reservoir for virulent disease (Nugues et al. 2004; Casey et al. 2014), microscale alterations of conditions in the benthic boundary layer (Carpenter and Williams 1993; Brown and Carpenter 2013), and physical interactions between polyps and algae (River and Edmunds 2001; Box and Mumby 2007). Wells et al. (2021) suggested that turf algae may negatively impact octocoral recruits by providing habitat to corallivorous mesofauna, although their observations were qualitative.

While octocorals and scleractinians are both negatively affected by algal competition (Lirman 2001; Hughes et al. 2007; Kuffner et al. 2006; Lasker and Kim 1996; Wells et al. 2021), they differ in their apparent response to grazers. Grazers frequently facilitate scleractinian growth, fecundity, and recruitment by removing algae (Lirman 2001; Stockton and Edmunds 2021; Nugues and Bak 2006; Foster, Box, and Mumby 2008; Idjadi, Haring, and Precht 2010). In some circumstances, urchin grazing can reduce recruitment and growth by direct consumption of coral tissue (Bak and van Eys 1975; O’Leary et al. 2013; Dang et al. 2020; Davies, Matz, and Vize 2013) and some species of particularly palatable coral are excluded from reef habitats by parrotfish through direct grazing (Littler, Taylor, and Littler 1989; Miller and Hay 1998). Fish reduce competition between recruits and macroalgae at the cost of incidental and targeted predation by fish. Doropoulos et al. (2016) has provided an in-depth analysis of this trade-off – scleractinian corals settled within crevices on exposed surfaces where fish removed algal competitors but could not consume the recruits. The impact of grazers on octocorals is less studied, but the available observations suggest grazing negatively affects octocoral recruitment. Octocoral recruitment increased two orders of magnitude in Puerto Rican reefs in 1984, concurrent with a mass mortality of the primary grazing sea urchin *Diadema antillarum* (Yoshioka and Yoshioka 1989; Yoshioka 1994). Grazing urchins also affected octocoral recruit distribution in the temperate Gulf of Maine; recruits were more often found among adult conspecifics where they survived longer than those that settled outside of adult aggregations (Sebens 1983b). In several recruitment studies of single polyp octocorals, there was evidence of grazing fish consuming octocoral recruits (Wells et al. 2021; Evans, Coffroth, and Lasker 2013), although in both studies, predation was never directly observed.

In this study, we used field experiments to quantify fish and urchin impacts on newly-settled octocoral polyps by excluding fish with cages and caging *D. antillarum* with octocoral polyps. We also quantified the impacts of mesofauna on recruitment in a laboratory experiment by naturally and experimentally manipulating the abundance of mesofauna. We hypothesized that grazers and mesofauna would negatively affect octocoral recruitment through incidental or intentional predation of primary polyps.

## Methods

The effects of grazing and predation on octocoral recruitment were assessed through a series of sequential experiments (Supp. Fig. 1) using unglazed terracotta settlement tiles (15 × 15 × 1 cm). Settlement tiles (*n* = 40) were initially deployed and conditioned in an octocoral-dominated reef in Grootpan Bay, St. John, U.S. Virgin Islands (18.309° N, 64.719° W) on April 12, 2021 for 62 days. This site is also known as East Cabritte in other studies (Lasker et al. 2020; Martínez-Quintana and Lasker 2021). The tile undersides had 22 3-mm deep vertical grooves providing refugia for algae and invertebrate recruits. Tiles were attached to the reef with 10-cm stainless steel rods with the refugia side 2-5 cm above the substratum.

**Figure 1.**
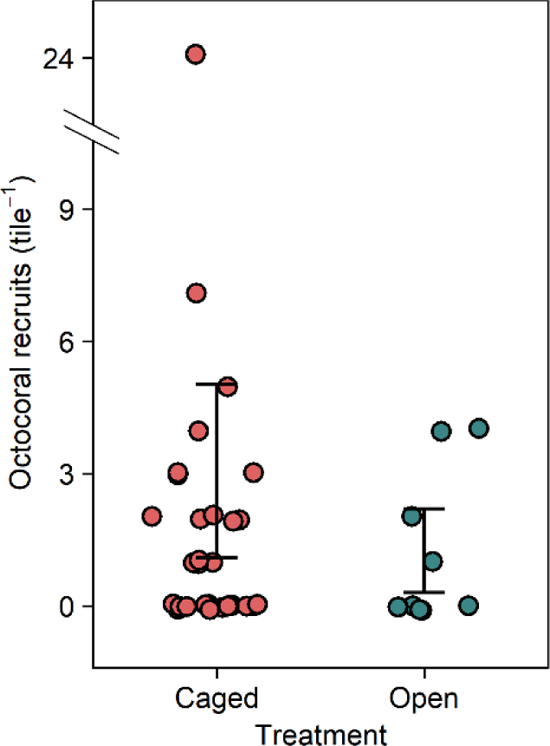
Effect of tile treatment on the number of recruited octocorals. Error bars are bootstrapped 95% confidence intervals of the means.

### Initial caging experiment

After conditioning, 30 of the tiles were caged in polyvinyl chloride-coated steel mesh (1.3 × 1.3 cm square mesh, 25 × 25 × 15 cm cages) to exclude fish from feeding on algae and invertebrates. Ten tiles were left open to fish grazing. We scrubbed the cages three weeks after deployment to remove fouling organisms and facilitate water movement. Tiles were collected after 47 days. We mapped the positions of any new octocoral recruits that settled while the tiles were on the reef so that we could identify individuals throughout subsequent experiments.

### Invertebrate removal experiment

Tiles from the initial caging experiment were brought into the laboratory to test the effect of invertebrate predation on octocoral recruitment. We applied one of three treatments to the previously caged tiles – tiles were either (1) left in a sea table with ten non-terminal bluehead wrasse *Thalassoma bifasciatum* (8-11 cm standard length, herein referred to as “fish removal” tiles), which removed most of the motile invertebrates; (2) soaked in 7.5% magnesium chloride for 30 minutes, shaken briefly to dislodge motile invertebrates, and then large solitary invertebrates were removed by hand (herein referred to as “manual removal” tiles); or (3) were left alone (herein referred to as “caged” tiles). The ten tiles left outside cages in the field were not manipulated and will be referred to as “open” tiles. The amphinomid worm *Hermodice carunculata*, where present, was removed from all treatments as it is a voracious predator of coral recruits (Wolf and Nugues 2013) and would likely hide the effect of other potential invertebrate predators. Invertebrates removed from three of the manual removal tiles were quantified to determine what motile invertebrate taxa were present. At the conclusion of the experiment, invertebrates from one tile of each treatment were identified and counted to determine the effectiveness of each invertebrate removal treatment by soaking them in magnesium chloride for 30 minutes, shaking them, and then manually picking any remaining invertebrates off the tile. The undersides of the tiles, where octocorals primarily settle (Wells et al. 2021; Martínez-Quintana and Lasker 2021), were photographed to categorize benthic cover after treatments were applied. Turf height was measured in five locations on the underside of the tiles.

Tiles were placed in 14 L clear plastic containers, one tile per container, refuge side down, and elevated by a ring of mesh as in Wells et al. (2021). Water circulation in each container was provided by bubbling air from a glass Pasteur pipette attached to an air pump. Water movement was slow but circulation was observed throughout the container. Ambient sunlight was provided from large south facing windows and the temperature of the room was held at 28 °C. 25% of the water in each container was changed daily. Octocoral planulae were collected from the water column within five meters of the surface at Grootpan Bay on August 1, 2021 (nine days after the full moon). Fifty-three competent octocoral planulae were added to each container (*n* = 36, nine containers per treatment). Planulae settled over the following ten days, after which all polyps on the tiles were mapped again. Polyps that were present before the experiment were identifiable by their previously mapped position and size relative to newly-settled polyps and were excluded from the analyses of the invertebrate removal experiment since they settled prior to the tiles being treated.

### Urchin inclusion and fish exclusion experiment

To assess the effects of urchin and fish grazing on octocoral survival and recruitment we redeployed the tiles onto steel rods in Grootpan Bay eleven days after we introduced planulae to the tiles in the laboratory experiment, with those tiles originally in cages placed back into cages to exclude fish. One small (1.5-2.5 cm test diameter) long-spined sea urchin *Diadema antillarum* was added to half of the cages to test the effect of urchin grazing on field settlement and survival. Settlement of exogenous octocoral recruits and survival of all octocoral polyps was tracked over the course of 18 days, with observations on days 0, 3, 6, 12, and 18. Photographs of the undersides of the tiles were taken on days 0, 6, 12, and 18 to correlate settlement and survival with benthic cover of invertebrates and algae.

### Benthic cover analyses

For all photographs of tiles throughout the experiments (laboratory and field experiments) benthic cover was quantified using CoralNet (Beijbom 2015). Initially, 25 randomly distributed points were overlayed on each image and benthic cover was manually annotated by CDW. These data (4250 annotations across 170 images) were used to train an automatic annotator using a computer vision algorithm (Beijbom et al. 2012). Each image was then overlayed with an 18 × 18 grid of points (324 points per image). These points were annotated with the vision algorithm. Automatic annotations that the vision algorithm identified with ≥84% confidence were 95% accurate in their identifications. Therefore, where the algorithm was <84% confident, points were subsequently manually annotated by CDW. A 5% reduction in annotation accuracy has marginal impacts on cover estimates (Beijbom et al. 2015). Benthic cover was identified to the following categories: bare space, colonial and solitary ascidians, bryozoans, crustose coralline algae, non-coralline encrusting algae, upright algae, sponges, turf algae, coral polyps, mollusk eggs, calcareous tubes and proteaceous tubes of worms. These broad categories were chosen to characterize general shifts in community composition on the tiles (e.g., increasing turf cover) without the destructive sampling that would be required to identify taxa at higher resolution.

### Statistical Analyses

The effects of initial fish exclusion via caging, invertebrate removal, and sea urchin inclusion/fish exclusion on octocoral recruitment were tested with generalized linear models (GLM) with Poisson error distributions in R version 4.2.1 (R Core Team 2022). Equidispersion was checked (R package AER 1.2-10, Kleiber and Zeileis 2008) and when models were not equidispersed, a negative binomial error distribution was used (R package MASS 7.3-57, Venables and Ripley 2002). These GLMs were run with treatment as a fixed factor. 95% bias-corrected and accelerated bootstrap confidence intervals were calculated with 100,000 replicates (R package bcaboot 0.2-3, Efron and Narasimhan 2021).

Benthic cover in each experiment was visualized with non-metric multidimensional scaling (nMDS) using a Bray-Curtis dissimilarity index with 9999 random starts to find a stable solution with two dimensions (R package vegan 2.6-2, Oksanen et al. 2022). When the final solution had a stress over 0.18, three dimensions were used. The effect of individual benthic cover types on recruitment were assessed for cover types with more than 1% of the total cover using smoothing functions in generalized additive models (GAMs). To determine the effect of turf height on octocoral recruitment in the invertebrate removal experiment, an additional GAM was run with turf height in a smoothing function and recruitment as the response variable. In the urchin inclusion and fish exclusion experiment, the effect of exposure to predation by fish and urchins on recruitment was analyzed using a GAM with sampling day as a smooth and treatment as a fixed factor. All GAMs were run (R package mgcv 1.8-40, Wood 2021). Variance components were estimated by restricted maximum likelihood with penalized cubic regression splines for smoothing, three knots for basis construction, and a negative binomial error distribution. Basis dimension choices were accessed for oversmoothing and where basis functions were too small, we added more knots, up to ten. Additionally, we tested for concurvity in smooths.

Time-dependent Cox proportional hazard models (CPH, Cox 1972) were used to determine the effect of exposure to urchins, fish, and benthic cover on survival of octocoral polyps (R package survival 3.3-1, Therneau 2022). The assumption that the relative effect of the hazard is proportional was tested prior to the analysis. When this assumption failed, piece-wise exponential additive mixed models (PAMM) were used (R package pammtools 0.5.8, Bender, Groll, and Scheipl 2018). PAMMs allow for time-varying covariates such as benthic cover and allow for the calculation of hazard ratios without the need for proportional hazards (Bender, Groll, and Scheipl 2018). Hazard ratios were calculated to compare treatments and levels of cover types with a significant effect on survival. Hazard ratios, in this case, depict the probability of mortality between time-steps for each treatment relative to the probability of mortality between time-steps for either a reference category for discrete variables or a reference value for continuous variables. For these models, we used either “caged” tiles as the reference category or median cover for benthic cover analyses. Hazard ratios significantly greater than 1.0 indicate that level is more hazardous than the reference, while those below 1.0 indicate it is less hazardous.

## Results

### Initial caging experiment

In the initial caging experiment 74 octocoral polyps recruited across the 40 tiles in 47 days. Recruitment was significantly higher on tiles within cages compared to uncaged tiles (Fig. 1, GLM, *p* = 0.02). Mean recruitment on caged tiles was 2.1 polyps × tile^−1^ (95% CI [1.2 – 5.1]) whereas open cages had a mean recruitment of 1.0 polyps × tile^−1^ (95% CI [0.2 – 2.0]). nMDS characterizations of composition on the undersides of the tiles greatly overlapped among treatments (Supp. Fig. 2). Octocoral recruitment was positively associated with increased levels of turf algae (Fig. 2A, GAM, *p* = 0.03) and negatively associated with increased levels of colonial ascidians (Fig. 2B, GAM, *p* < 0.01).

**Figure 2.**
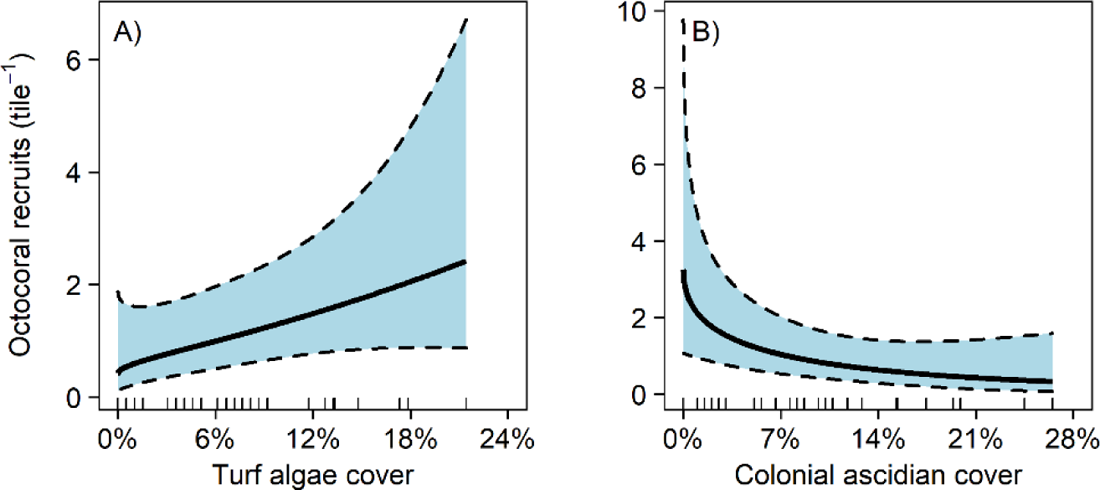
Effect of turf algae and colonial ascidian cover on octocoral recruitment on the underside of settlement tiles in the initial caging experiment. Bold lines are fitted predictions from a generalized additive model with 95% confidence intervals. Predictor variables are shown by rug plots on each abscissa.

### Invertebrate removal experiment

In the invertebrate removal experiment 606 of 1908 octocoral planulae (32%) recruited across the 36 tiles. Octocoral recruitment did not vary significantly between removal treatments (Fig. 3, GLM, *p* = 0.71). Mean recruitment across all treatments was 5.8 polyps tile^−1^ (95% CI [4.9 – 6.7]).

**Figure 3.**
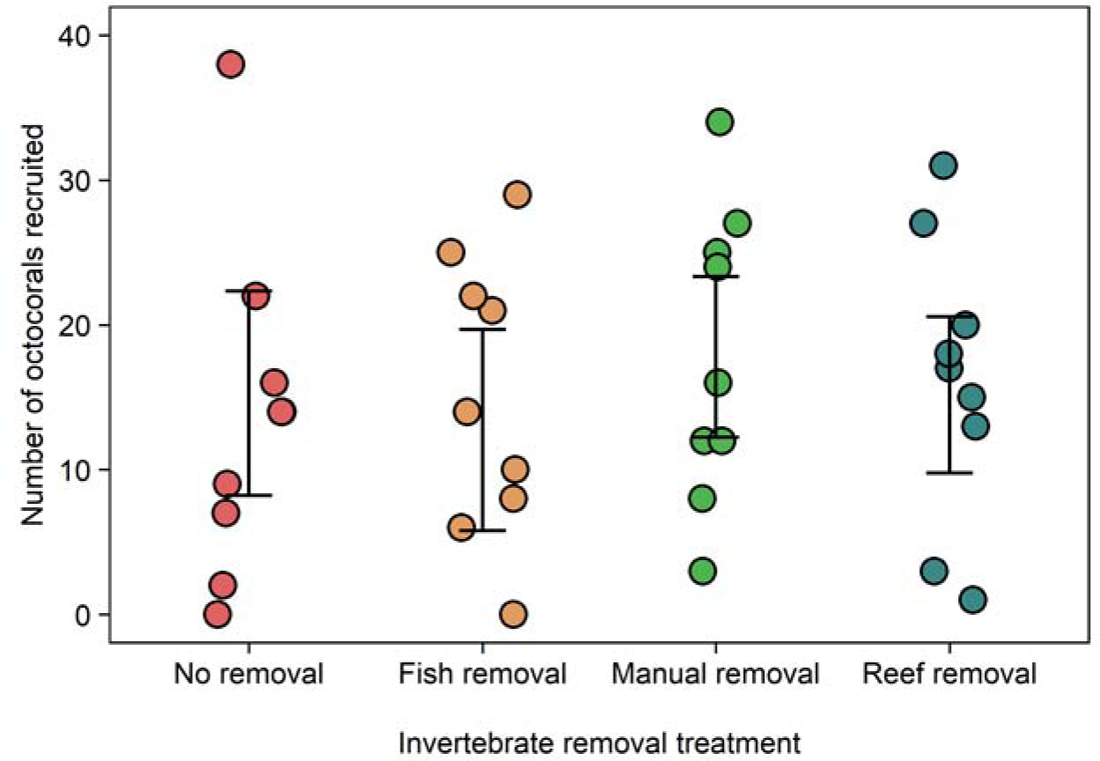
Effect of invertebrate removal treatment on the number of recruited octocorals. Error bars are bootstrapped 95% confidence intervals of the means.

The manual removal treatment was effective at removing annelid worms, arthropods, mollusks, and nematodes (Supp. Table 1). Throughout the experiment, invertebrate abundance decreased across all treatments except for copepods, cephalocaridians, and nematodes in the fish removal treatment; nematodes in the manual removal treatment; and isopods in the reef removal treatment (Supp. Table 2). Bare space on the tiles became more abundant over the course of the experiment although tiles that were originally caged in the initial caging experiment maintained higher levels of turf throughout the laboratory experiment (Supp. Fig. 3). Octocoral recruitment did not significantly respond to any individual benthic cover category (GAM, *p* ≥ 0.12) nor any dimensions of the nMDS (GAM, *p* ≥ 0.09). Octocoral recruitment did not significantly respond to turf height (GAM, *p* = 0.12).

### Urchin inclusion and fish exclusion experiment

There was a total of 587 polyps on the 33 tiles outplanted from the laboratory settlement experiment to the reef. After 18 days, 312 (53%) of those polyps survived. The urchin inclusion and non-caged treatments were significantly more hazardous to octocoral polyps than the caged treatment (CPH, *p* ≤ 0.03) with a 35% higher chance of dying in the next time step when exposed to an urchin and a 67% higher chance of dying in the next time step when exposed to fish (Fig. 4A). Increased cover of coralline algae, calcified worm tubes, and turf algae were associated with reduced survival of octocoral polyps (PAMM, *p* ≤ 0.04). For every 1% increase in coralline algae, calcified worm tubes, and turf algae cover, there was a respective 4.0%, 2.5%, and 1.7% increase in the likelihood of dying in the next time step.

**Figure 4.**
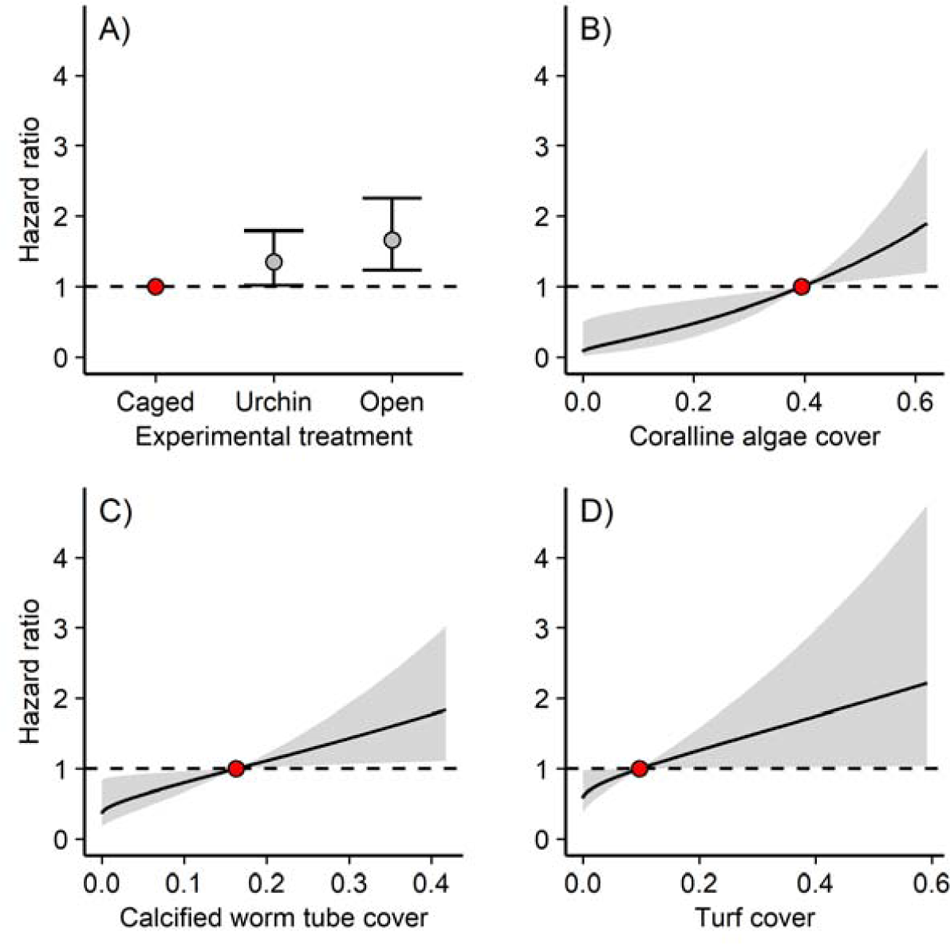
Effect of (A) experimental treatment, (B) coralline algae cover, (C) calcified worm tube cover, and (D) turf cover on hazard ratio. The hazard ratio is the probability of mortality in the next time step relative to the probability of mortality of the reference value (the red point). Reference values for benthic cover types is the median cover. Error bars and shaded areas are 95% confidence intervals of the hazard ratio.

In the urchin inclusion and fish exclusion experiment, 191 polyps recruited across the 33 tiles in 18 days. Octocoral recruitment did not have a significant response to the invertebrate removal treatments (Fig. 5A, GLM, *p* = 0.71) but was significantly related to sample day, with more recruitment at the beginning and end of the experiment than in the middle (Fig. 5B, GAM, *p* < 0.01). High settlement occurred during periods when many planulae were observed in the water column. Mean recruitment across all treatments was 5.8 polyps × tile^−1^ (95% CI [4.9 – 6.7]). nMDS characterizations of composition on the undersides of the tile greatly overlapped among treatments and did not change greatly over time (Supp. Fig 4). Octocoral recruitment on the undersides of the tiles was negatively associated with increased levels of coralline algae (Fig. 6, GAM, *p* = 0.02).

**Figure 5.**
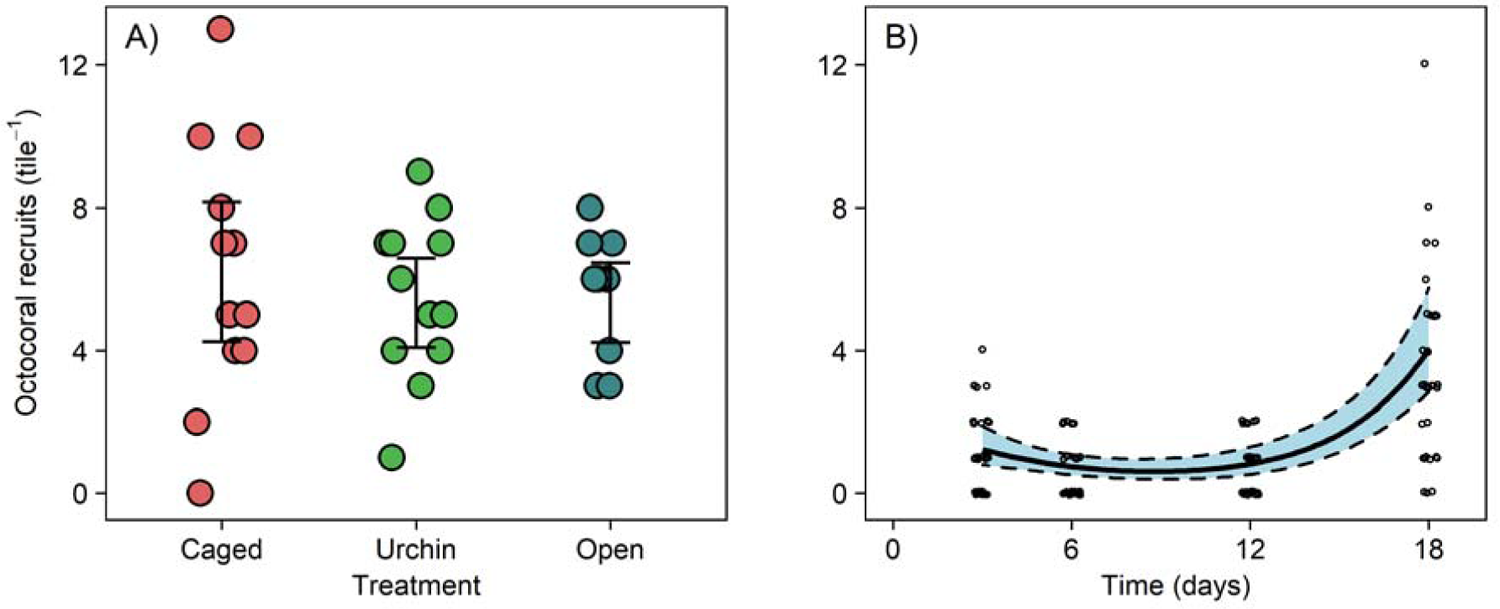
Effect of (A) experimental treatment and (B) sampling day on exogenous octocoral settlement. Error bars are bootstrapped 95% confidence intervals of the means. The bold line is the fitted prediction from a generalized additive model with shaded 95% confidence intervals. Data points are jittered along both axes for visibility.

**Figure 6.**
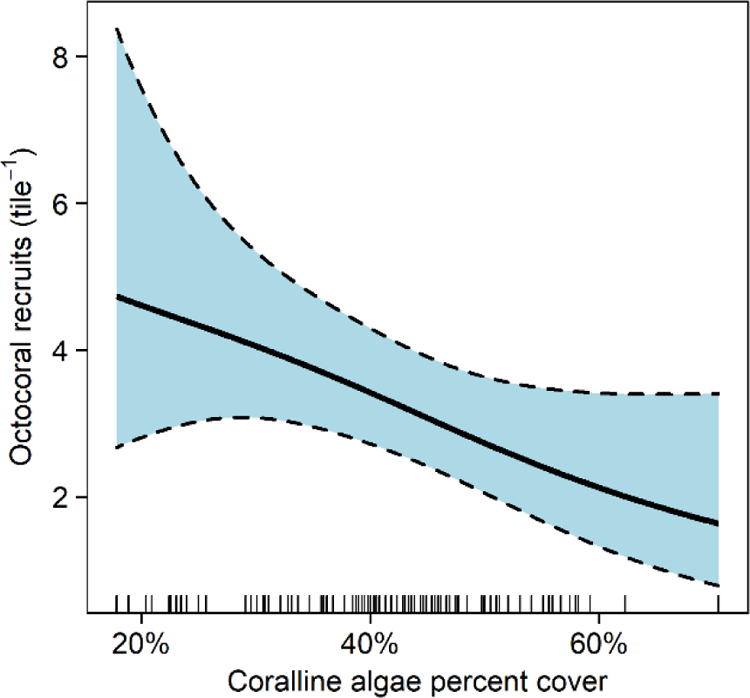
Effect of coralline algae cover on octocoral settlement in the urchin inclusion and fish exclusion experiment. The bold line is a fitted prediction from a generalized additive model with shaded 95% confidence intervals. Predictor variables are shown by a rug plot on the abscissa.

## Discussion

Benthic grazers on coral reefs can facilitate the growth of corals by grazing on their competitors (e.g., turf algae) or hinder corals through predation of coral tissue and recruits. In our experiments, exposure to fish and urchin grazing significantly reduced survival of single-polyp octocorals (Fig. 1 and Fig. 4A), while mesofauna did not significantly affect recruitment (Fig. 3). We also found significant relationships between recruitment and benthic cover in our field experiments (Fig. 2, 4, and 5), but these correlations likely reflect exposure to grazing.

### Exposure to urchin grazing

As expected, exposure to *D. antillarum* urchin grazing led to decreased survival (Fig. 4A). This result supports the findings of Yoshioka and Yoshioka (1989), where recruitment of octocorals increased by two orders of magnitude on Puerto Rican reefs after the catastrophic die-off of *D. antillarum* in December of 1984. Within urchin inclusion cages, sections of the tiles were denuded of live cover, except for coralline crusts and calcium carbonate worm tubes. Octocoral recruits that had settled within those areas were all removed. It is unknown whether removal was via direct consumption or via incidental dislodgement while the urchin grazed on the benthic community although the effect is the same.

Despite the negative effects of urchins on survival, octocorals did not avoid settling on tiles with caged urchins (Fig. 5A). Settlement was similar to settlement levels on tiles in cages without urchins or tiles outside of cages. Settlement in the field was only related to the sampling day (Fig 4B). We observed elevated settlement approximately ten days after the full moon, which is approximately five days after predicted spawning for a number of octocoral species (Kahng, Benayahu, and Lasker 2011), indicating that octocorals prefer to settle as soon as they are competent. This timing matches the findings of Tonra, Wells, and Lasker (2021) and Lasker and Kim (1996), where Caribbean plexaurid octocorals took five days for median competency. Plexaurids are the most abundant family of octocorals that spawned during the time of this study (Kahng, Benayahu, and Lasker 2011). The highest settlement pulse was coincident with times when there were many planulae in the water column similar to observations by Wells et al. (2021).

Similar experiments examining the interactions between urchins and scleractinian corals have found the relationship between the two taxa to be facilitatory in some circumstances and detrimental in others for scleractinians. Generally, urchin grazing improves recruitment of scleractinians by removing macroalgae and exposing crustose coralline algae (Edmunds and Carpenter 2001; Carpenter and Edmunds 2006; Nozawa, Lin, and Meng 2020). Urchin grazing capabilities should scale with urchin size. Indeed, small urchins (1 cm *Mespilia globulus*), in an urchin-coral co-culturing experiment, improved growth and survival of scleractinians by reducing algal competition, even at high urchin densities (75 m^−2^), so high that urchin growth was slowed by intraspecific competition for algae (Craggs et al. 2019). However, urchins can reduce recruitment of particularly palatable scleractinian species on surfaces without grazing refugia and when urchins are in high density (Sammarco 1980; Davies, Matz, and Vize 2013; Dang et al. 2020). In experiments where adult urchin densities were manipulated, coral survival was highest at intermediate levels of urchin density. When urchin densities were too high, urchins directly injured or killed scleractinians and when urchin densities were too low, scleractinians were outcompeted by algae. Sammarco (1980) found that 16 adult urchins m^−2^ reduced scleractinian recruitment by an order of magnitude compared to replicates without *D. antillarum*. We had one small *D. antillarum* (≤ 2.5 cm test diameter) in each replicate (density = 16 m^−2^) with 3 mm deep crevices as potential refugia. This density of urchins was too high or the refugia were insufficient to protect octocorals from grazing pressure.

Prior to the catastrophic die-off of *D. antillarum*, octocoral recruitment was likely reduced by urchins and was subsequently released from this pressure after the die-off, leading to the impressive recruitment pulse observed by Yoshioka and Yoshioka (1989). In the decades since the first die-off, *D. antillarum* recovery has been modest with the densest populations in the east Caribbean (reviewed in Lessios 2016), which may restore some of the ecosystem services provided by *D. antillarum*. Additionally, there have been small-scale experimental restoration efforts to bring back *D. antillarum* (Maciá, Robinson, and Nalevanko 2007; Williams 2022). However, a wave of *D. antillarum* die-off is currently occurring across the Caribbean (Atlantic and Gulf Rapid Reef Assessment 2022). Unlike the 1983-1984 die-off where octocoral recruitment was released from mortality from grazing *D. antillarum* (Yoshioka and Yoshioka 1989; Yoshioka 1996), another die-off is unlikely to improve octocoral recruitment because urchin populations are long way from fully recovering.

### Exposure to fish predation and grazing

Similar to the effect of urchins, exposure to fish decreased survival, but not settlement, of octocorals (Fig. 1, 4A, and 5). While predation was never directly observed, fourspot butterfly fish (*Chaetodon capistratus*) were frequently seen inspecting the open tiles, presumably looking for cnidarian polyps and annelids, their preferred foods (Liedke et al. 2018; Lasker 1985). In addition, parrotfish (scarids) and tangs (acanthurids) were observed grazing on the tiles and grazing scars from fish were abundant, especially on the upper surfaces, similar to observations made by Wells et al. (2021). In a similar fish exclusion study, octocoral survival was significantly improved within cages where fish predation was reduced (Evans, Coffroth, and Lasker 2013). In the same study, scleractinian recruit survival and fish exposure were not related. Scleractinian recruits seem to be avoided by grazing fishes (Birkeland 1977; Brock 1979), although some grazing fish with relatively large gapes and less discriminating grazing behavior can remove recruits (Christiansen et al. 2009). In some locations, reduced scleractinian recruitment has been associated with elevated grazing pressure (Penin et al. 2010). Scleractinian recruits are partially protected from predation by their calcium carbonate skeleton, which octocorals lack. Many adult octocorals employ secondary metabolites and/or abundant calcium carbonate sclerites as an antipredator deterrent, but regardless of their defenses, if a grazer inadvertently consumed an octocoral recruit and rejected it as food, the result is the same and the recruit dies.

### Exposure to mesofauna

Octocoral recruitment was not reduced by mesofauna abundance in the laboratory experiment (Fig. 3). Wells et al. (2021) hypothesized that the reduced survival in turf algae was due to the elevated levels of mesofauna within the turf. Our results demonstrate that mesofauna do not affect survival. Other potential hypotheses that have been supported with scleractinians include allelopathic competition, algae providing a reservoir for virulent diseases, microscale alterations of conditions in the benthic boundary layer, and physical interactions between polyps and upright algae (Kuffner et al. 2006; Rasher et al. 2011; Bonaldo and Hay 2014; Nugues et al. 2004; Casey et al. 2014; Carpenter and Williams 1993; Brown and Carpenter 2013; River and Edmunds 2001; Box and Mumby 2007). While mesofauna did not reduce recruitment, there are several invertebrates that actively consume octocorals such as the amphinomid worm *H. carunculata* and ovulid gastropods in the genus *Cyphoma* (Vreeland and Lasker 1989; Lasker and Coffroth 1988; Burkepile and Hay 2007). Both of these taxa were excluded from the laboratory experiments. Caged treatments in the field did not exclude *H. carunculata*, but should have excluded larger *Cyphoma* spp. Neither taxa has been documented consuming single polyp recruits although both have been observed eating less than 5-cm tall colonies (Edmunds and Lasker 2019). It seems that fish predation and urchin grazing are more important factors to recruit survival than predation by mesofauna for primary polyps.

### Effect of benthic cover

In the initial caging experiment, which excluded fish predation and grazing as well as large invertebrates, octocorals recruited significantly more within cages (Fig. 1). Regardless of caging treatment, tiles with more turf algae on their undersides had higher octocoral recruitment (Fig. 2A). Conversely, octocoral recruitment was reduced on tiles with more colonial ascidians (Fig. 2B). Neither cover category dominated the undersides. Both were less than 9% of the total benthic cover and there was sufficient empty tile space for octocoral settlement (11%). In addition, there was an average of 35% crustose coralline algae cover, a preferred settlement location for some octocorals (Lasker and Kim 1996; Slattery et al. 1999). One hypothesis for this relationship is that these two benthic groups, turf algae and colonial ascidians, were indicator taxa for how grazed the underside of a tile was with more grazed tiles having less turf and more colonial ascidians. Interestingly, Wells et al. (2021) found that more octocorals recruited and survived longer on tiles with less turf algae, opposite of this study. Perhaps, at low levels of turf cover, the competition between octocoral recruits and turf algae is less detrimental to octocoral recruitment than the presence of potential grazers and so both taxa increase in abundance together controlled by a similar mechanism. We observed several instances of sponges and colonial ascidians overgrowing octocoral recruits, but generally these interactions were limited to a few recruits on a few tiles that settled directly next to these taxa.

Unexpectedly, in the urchin inclusion and fish exclusion experiment, tiles with more crustose coralline algae had less settlement (Fig. 6) and tiles with more coralline algae were more hazardous to recruits (Fig. 4B). That result is in contrast to many studies indicating the importance of coralline algae on settlement of octocorals (Lasker and Kim 1996; Slattery et al. 1999; Sebens 1983a; Benayahu, Achituv, and Berner 1989). However, not all corals are induced to settle by coralline algae and not all coralline algae induce settlement of corals (Morse et al. 1988; Heyward and Negri 1999). Additionally, coralline algae can actively deter settlement and defend themselves from being fouled by shedding epithelial cells, overgrowing organisms that grow on them, and exuding allelopathic chemicals (Keats, Knight, and Pueschel 1997; Sebens 1986; Harrington et al. 2004). Thus, the inverse relationship between successful recruitment and coralline algal cover could represent an interaction between the two taxa. Alternatively, the abundant coralline algae, like turf algae and colonial ascidians in the initial caging experiment, may be a product of heavier grazing pressure which would reduce both survival of the polyps originally on the tiles and *in situ* recruitment after the tiles were outplanted on the reef. The negative relationship between polyp survival and calcified worm tubes (Fig. 4C) indicates that this might be the most likely explanation as there are no studies that indicate calcified worm tubes should negatively affect coral recruits. In fact, some cnidarians can be induced to settle when presented with calcified worm tubes (Chia and Spaulding 1972). Calcified worm tubes were present in similar abundances on all tiles, but they were only visible in the images used to determine abundance when they were not under other benthic cover groups such as turf algae. Opposite to the initial caging experiment but similar to findings of Wells et al. (2021), turf algae reduced survival in the urchin inclusion and fish exclusion experiment (Fig. 4D). This disparity may be due the difference in ranges of turf cover in the two experiments. In the initial caging experiment, turf algae were always less than 21% cover whereas in the urchin inclusion and fish exclusion experiment the turf cover ranged from 0 to 59%. At higher levels of turf cover, competition with turf algae may become more detrimental, reducing survival.

### Summary

Caribbean octocoral recruits are sensitive to urchin and fish grazing, but invertebrate mesofauna predation, excluding ovulid snails and amphinomid worms, seems to be a less important factor. Scleractinian growth, fecundity, and recruitment are frequently facilitated by grazers (Lirman 2001; Stockton and Edmunds 2021; Nugues and Bak 2006; Foster, Box, and Mumby 2008; Idjadi, Haring, and Precht 2010), opposite to octocorals, despite them both being sensitive to algal competition (Lirman 2001; Hughes et al. 2007; Kuffner et al. 2006; Lasker and Kim 1996; Wells et al. 2021). Unlike scleractinians, octocorals lack a protective skeleton which protects at least some of the tissue from complete predation by grazers and although they produce secondary metabolites, which reduce predation on adult octocorals, chemical defense seems ineffective at reducing predation of recruits. The removal of large herbivorous fish and the die-off of the urchin *D. antillarum* may have led to increased recruitment of octocorals and contributed to the increase in abundance of octocorals in the Caribbean, concurrent with the loss of scleractinians.

## Acknowledgements

This work was completed under a permit from the Virgin Islands National Park (VIIS-2021-SCI-0008) and was funded by the National Science Foundation (OCE 17-56381). We thank the Virgin Islands Environmental Research Station and the University of the Virgin Islands for laboratory space and P.J. Edmunds for reading and commenting on the manuscript.

## References

Atlantic and Gulf Rapid Reef Assessment. 2022. “Diadema response network.” Accessed July 25. https://www.agrra.org/sea-urchin-die-off/.

Bak, R. P. M., and G. van Eys. 1975. “Predation of the sea urchin Diadema antillarum Philippi on living coral.” Oecologia 20 (2): 111–115. https://doi.org/10.1007/BF00369023.

Bartlett, L. A., V. I. P. Brinkhuis, R. R. Ruzicka, M. A. Colella, K. S. Lunz, E. H. Leone, and P. Hallock. 2018. “Chapter 6. Dynamics of stony coral and octocoral juvenile assemblages following disturbance on patch reefs of the Florida Reef Tract.” In Corals in a changing world, edited by Carmenza Duque and Edisson Tello Camacho, 99-120. London, UK: IntechOpen.

Beijbom, O. 2015. “Automated annotation of coral reef survey images.” Doctor of Philosophy Dissertation, Computer Science and Engineering, University of California, San Diego.

Beijbom, O., P. J. Edmunds, D. I. Kline, B. G. Mitchell, and D. Kreigman. 2012. “Automated annotation of coral reef survey images.” 2012 IEEE Conference on Computer Vision and Pattern Recognition, 16-21 June 2012.

Beijbom, O., P. J. Edmunds, C. Roelfsema, J. Smith, D. I. Kline, B. P. Neal, M. J. Dunlap, V. Moriarty, T.-Y. Fan, C.-J. Tan, S. Chan, T. Treibitz, A. Gamst, B. G. Mitchell, and D. Kriegman. 2015. “Towards automated annotation of benthic survey images: Variability of human experts and operational modes of automation.” PLoS One 10 (7): e0130312. https://doi.org/10.1371/journal.pone.0130312.

Bellwood, D. R., T. P. Hughes, C. Folke, and M. Nyström. 2004. “Confronting the coral reef crisis.” Nature 429 (6994): 827–833. https://doi.org/10.1038/nature02691.

Benayahu, Y., Y. Achituv, and T. Berner. 1989. “Metamorphosis of an octocoral primary polyp and its infection by algal symbiosis.” Symbiosis 7: 159–169.

Bender, A., A. Groll, and F. Scheipl. 2018. “A generalized additive model approach to time-to-event analysis.” Statistical Modelling 18 (3-4): 299–321. https://doi.org/10.1177/1471082X17748083.

Birkeland, C. 1977. “The importance of rate of biomass accumulation in early successional stages of benthic communities to the survival of coral recruits.” Third International Coral Reef Symposium, Miami, Florida, US.

Bonaldo, R. M., and M. E. Hay. 2014. “Seaweed-coral interactions: Variance in seaweed allelopathy, coral susceptibility, and potential effects on coral resilience.” PLoS One 9 (1): e85786. https://doi.org/10.1371/journal.pone.0085786.

Box, S. J., and P. J. Mumby. 2007. “Effect of macroalgal competition on growth and survival of juvenile Caribbean corals.” Marine Ecology Progress Series 342: 139–149. https://doi.org/10.3354/meps342139.

Brock, R. E. 1979. “An experimental study on the effects of grazing by parrotfishes and role of refuges in benthic community structure.” Marine Biology 51 (4): 381–388. https://doi.org/10.1007/BF00389216.

Brown, A. L., and R. C. Carpenter. 2013. “Water-flow mediated oxygen dynamics within massive Porites-algal turf interactions.” Marine Ecology Progress Series 490: 1–10. https://doi.org/10.3354/meps10467.

Bruno, J. F., H. Sweatman, W. F. Precht, E. R. Selig, and V. G. W. Schutte. 2009. “Assessing evidence of phase shifts from coral to macroalgal dominance on coral reefs.” Ecology 90 (6): 1478–1484. https://doi.org/10.1890/08-1781.1.

Burkepile, D. E., and M. E. Hay. 2007. “Predator release of the gastropod Cyphoma gibbosum increases predation on gorgonian corals.” Oecologia 154 (1): 167–173. https://doi.org/10.1007/s00442-007-0801-4.

Carpenter, R. C., and P. J. Edmunds. 2006. “Local and regional scale recovery of Diadema promotes recruitment of scleractinian corals.” Ecology Letters 9 (3): 271–280. https://doi.org/10.1111/j.1461-0248.2005.00866.x.

Carpenter, R. C., and S. L. Williams. 1993. “Effects of algal turf canopy height and microscale substratum topography on profiles of flow speed in a coral forereef environment.” Limnology and Oceanography 38 (3): 687–694. https://doi.org/10.4319/lo.1993.38.3.0687.

Casey, J. M., T. D. Ainsworth, J. H. Choat, and S. R. Connolly. 2014. “Farming behaviour of reef fishes increases the prevalence of coral disease associated microbes and black band disease.” Proceedings of the Royal Society B: Biological Sciences 281 (1788). https://doi.org/10.1098/rspb.2014.1032.

Chia, F.-S., and J. G. Spaulding. 1972. “Development and juvenile growth of the sea anemone, Tealia crassicornis.” Biological Bulletin 142 (2): 206–218. https://doi.org/10.2307/1540225.

Christiansen, N. A., S. Ward, S. Harii, and I. R. Tibbetts. 2009. “Grazing by a small fish affects the early stages of a post-settlement stony coral.” Coral Reefs 28 (1): 47–51. https://doi.org/10.1007/s00338-008-0429-9.

Cox, D. R. 1972. “Regression models and life tables.” Journal of the Royal Statistical Society. Series B (Methodological) 34 (2): 187–220. https://doi.org/10.1111/j.2517-6161.1972.tb00899.x.

Craggs, J., J. Guest, M. Bulling, and M. Sweet. 2019. “Ex situ co culturing of the sea urchin, Mespilia globulus and the coral Acropora millepora enhances early post-settlement survivorship.” Scientific Reports 9 (1): 12984. https://doi.org/10.1038/s41598-019-49447-9.

Dang, V. D. H., C.-L. Fong, J.-H. Shiu, and Y. Nozawa. 2020. “Grazing effects of sea urchin Diadema savignyi on algal abundance and coral recruitment processes.” Scientific Reports 10 (1): 20346. https://doi.org/10.1038/s41598-020-77494-0.

Davies, S. W., M. V. Matz, and P. D. Vize. 2013. “Ecological complexity of coral recruitment processes: Effects of invertebrate herbivores on coral recruitment and growth depends upon substratum properties and coral species.” PLoS One 8 (9): e72830–e72830. https://doi.org/10.1371/journal.pone.0072830.

Doropoulos, C., G. Roff, Y.-M. Bozec, M. Zupan, J. Werminghausen, and P. J. Mumby. 2016. “Characterizing the ecological trade-offs throughout the early ontogeny of coral recruitment.” Ecological Monographs 86 (1): 20–44. https://doi.org/10.1890/15-0668.1.

Edmunds, P. J., and R. C. Carpenter. 2001. “Recovery of Diadema antillarum reduces macroalgal cover and increases abundance of juvenile corals on a Caribbean reef.” Proceedings of the National Academy of Sciences 98 (9): 5067–5071. https://doi.org/10.1073/pnas.071524598.

Edmunds, P. J., and H. R. Lasker. 2016. “Cryptic regime shift in benthic community structure on shallow reefs in St. John, US Virgin Islands.” Marine Ecology Progress Series 559: 1–12. https://doi.org/10.3354/meps11900.

Edmunds, P. J. 2019. “Regulation of population size of arborescent octocorals on shallow Caribbean reefs.” Marine Ecology Progress Series 615: 1–14. https://doi.org/10.3354/meps12907.

Efron, B., and B. Narasimhan. 2021. bcaboot: Bias corrected bootstrap confidence intervals R package version 0.2-3.

Evans, M. J., M. A. Coffroth, and H. R. Lasker. 2013. “Effects of predator exclusion on recruit survivorship in an octocoral (Briareum asbestinum) and a scleractinian coral (Porites astreoides).” Coral Reefs 32 (2): 597–601. https://doi.org/10.1007/s00338-012-1001-1.

Foster, N. L., S. J. Box, and P. J. Mumby. 2008. “Competitive effects of macroalgae on the fecundity of the reef-building coral Montastraea annularis.” Marine Ecology Progress Series 367: 143–152. https://doi.org/10.3354/meps07594.

Gardner, T. A., I. M. Côté, J. A. Gill, A. Grant, and A. R. Watkinson. 2003. “Long-term region-wide declines in Caribbean corals.” Science 301 (5635): 958–960. https://doi.org/10.1126/science.1086050.

Harrington, L., K. Fabricius, G. De’ath, and A. Negri. 2004. “Recognition and selection of settlement substrata determine post-settlement survival in corals.” Ecology 85 (12): 3428–3437. https://doi.org/10.1890/04-0298.

Heyward, A. J., and A. P. Negri. 1999. “Natural inducers for coral larval metamorphosis.” Coral Reefs 18 (3): 273–279. https://doi.org/10.1007/s003380050193.

Hughes, T. P. 1994. “Catastrophes, phase shifts, and large-scale degradation of a Caribbean coral reef.” Science 265 (5178): 1547–1551. https://doi.org/10.1126/science.265.5178.1547.

Hughes, T. P., K. D. Anderson, S. R. Connolly, S. F. Heron, J. T. Kerry, J. M. Lough, A. H. Baird, J. K. Baum, M. L. Berumen, T. C. Bridge, D. C. Claar, C. M. Eakin, J. P. Gilmour, N. A. J. Graham, H. Harrison, J.-P. A. Hobbs, A. S. Hoey, M. Hoogenboom, R. J. Lowe, M. T. McCulloch, J. M. Pandolfi, M. Pratchett, V. Schoepf, G. Torda, and S. K. Wilson. 2018. “Spatial and temporal patterns of mass bleaching of corals in the Anthropocene.” Science 359 (6371): 80–83. https://doi.org/10.1126/science.aan8048.

Hughes, T. P., M. J. Rodrigues, D. R. Bellwood, D. Ceccarelli, O. Hoegh-Guldberg, L. McCook, N. Moltschaniwskyj, M. S. Pratchett, R. S. Steneck, and B. Willis. 2007. “Phase shifts, herbivory, and the resilience of coral reefs to climate change.” Current Biology 17 (4): 360–365. https://doi.org/10.1016/j.cub.2006.12.049.

Idjadi, J. A., R. N. Haring, and W. F. Precht. 2010. “Recovery of the sea urchin Diadema antillarum promotes scleractinian coral growth and survivorship on shallow Jamaican reefs.” Marine Ecology Progress Series 403: 91–100. https://doi.org/10.3354/meps08463.

Kahng, S. E., Y. Benayahu, and H. R. Lasker. 2011. “Sexual reproduction in octocorals.” Marine Ecology Progress Series 443: 265–283. https://doi.org/10.3354/meps09414.

Keats, D. W., M. A. Knight, and C. M. Pueschel. 1997. “Antifouling effects of epithallial shedding in three crustose coralline algae (Rhodophyta, Coralinales) on a coral reef.” Journal of Experimental Marine Biology and Ecology 213 (2): 281–293. https://doi.org/10.1016/S0022-0981(96)02771-2.

Kleiber, C., and A. Zeileis. 2008. Applied econometrics in R R package version 1.2-9.

Kuffner, I. B., L. J. Walters, M. A. Becerro, V. J. Paul, R. Ritson-Williams, and K. S. Beach. 2006. “Inhibition of coral recruitment by macroalgae and cyanobacteria.” Marine Ecology Progress Series 323: 107–117. https://doi.org/10.3354/meps323107.

Lasker, H. R. 1985. “Prey preferences and browsing pressure of the butterflyfish Chaetodon capistratus on Caribbean gorgonians.” Marine Ecology Progress Series 21 (3): 213–220. https://doi.org/10.3354/meps021213.

Lasker, H. R., and M. A. Coffroth. 1988. “Temporal and spatial variability among grazers: Variability in the distribution of the gastropod Cyphoma gibbosum on octocorals.” Marine Ecology Progress Series 43 (3): 285–295. https://doi.org/10.3354/meps043285.

Lasker, H. R., and K. Kim. 1996. “Larval development and settlement behavior of the gorgonian coral Plexaura kuna (Lasker, Kim and Coffroth).” Journal of Experimental Marine Biology and Ecology 207 (1): 161–175. https://doi.org/10.1016/S0022-0981(96)02625-1.

Lasker, H. R., Á. Martínez-Quintana, L. Bramanti, and P. J. Edmunds. 2020. “Resilience of octocoral forests to catastrophic storms.” Scientific Reports 10 (1): 4286. https://doi.org/10.1038/s41598-020-61238-1.

Lenz, E. A., L. Bramanti, H. R. Lasker, and P. J. Edmunds. 2015. “Long-term variation of octocoral populations in St. John, US Virgin Islands.” Coral Reefs 34 (4): 1099–1109. https://doi.org/10.1007/s00338-015-1315-x.

Lessios, H. A. 2016. “The great Diadema antillarum die-off: 30 years later.” Annual Review of Marine Science 8 (1): 267–283. https://doi.org/10.1146/annurev-marine-122414-033857.

Liedke, A. M. R., R. M. Bonaldo, B. Segal, C. E. L. Ferreira, L. T. Nunes, A. P. Burigo, S. Buck, L. G. R. Oliveira-Santos, and S. R. Floeter. 2018. “Resource partitioning by two syntopic sister species of butterflyfish (Chaetodontidae).” Journal of the Marine Biological Association of the United Kingdom 98 (7): 1767–1773. https://doi.org/10.1017/S0025315417001321.

Linares, C., E. Cebrian, and R. Coma. 2012. “Effects of turf algae on recruitment and juvenile survival of gorgonian corals.” Marine Ecology Progress Series 452: 81–88. https://doi.org/10.3354/meps09586.

Lirman, D. 2001. “Competition between macroalgae and corals: effects of herbivore exclusion and increased algal biomass on coral survivorship and growth.” Coral Reefs 19 (4): 392–399. https://doi.org/10.1007/s003380000125.

Littler, M. M., P. R. Taylor, and D. S. Littler. 1989. “Complex interactions in the control of coral zonation on a Caribbean reef flat.” Oecologia 80 (3): 331–340. https://doi.org/10.1007/BF00379034.

Maciá, S., M. P. Robinson, and A. Nalevanko. 2007. “Experimental dispersal of recovering Diadema antillarum increases grazing intensity and reduces macroalgal abundance on a coral reef.” Marine Ecology Progress Series 348: 173–182. https://doi.org/10.3354/meps06962.

Martínez-Quintana, Á., and H. R. Lasker. 2021. “Early life-history dynamics of Caribbean octocorals: The critical role of larval supply and partial mortality.” Frontiers in Marine Science 8: 705563. https://doi.org/10.3389/fmars.2021.705563.

Miller, M. W., and M. E. Hay. 1998. “Effects of fish predation and seaweed competition on the survival and growth of corals.” Oecologia 113 (2): 231–238. https://doi.org/10.1007/s004420050373.

Morse, D. E., N. Hooker, A. N. C. Morse, and R. A. Jensen. 1988. “Control of larval metamorphosis and recruitment in sympatric agariciid corals.” Journal of Experimental Marine Biology and Ecology 116 (3): 193–217. https://doi.org/10.1016/0022-0981(88)90027-5.

Norström, A. V., M. Nyström, J. Lokrantz, and C. Folke. 2009. “Alternative states on coral reefs: Beyond coral–macroalgal phase shifts.” Marine Ecology Progress Series 376: 295–306. https://doi.org/10.3354/meps07815.

Nozawa, Y., C.-H. Lin, and P.-J. Meng. 2020. “Sea urchins (diadematids) promote coral recovery via recruitment on Taiwanese reefs.” Coral Reefs 39 (4): 1199–1207. https://doi.org/10.1007/s00338-020-01955-1.

Nugues, M. M., and R. P. M. Bak. 2006. “Differential competitive abilities between Caribbean coral species and a brown alga: A year of experiments and a long-term perspective.” Marine Ecology Progress Series 315: 75–86. https://doi.org/10.3354/meps315075.

Nugues, M. M., G. W. Smith, R. J. van Hooidonk, M. I. Seabra, and R. P. M. Bak. 2004. “Algal contact as a trigger for coral disease.” Ecology Letters 7 (10): 919–923. https://doi.org/10.1111/j.1461-0248.2004.00651.x.

O’Leary, J. K., D. Potts, K. M. Schoenrock, and T. R. McClahanan. 2013. “Fish and sea urchin grazing opens settlement space equally but urchins reduce survival of coral recruits.” Marine Ecology Progress Series 493: 165–177. https://doi.org/10.3354/meps10510.

Oksanen, J., G. L. Simpson, F. G. Blanchet, R. Kindt, P. Legendre, P. R. Minchin, R. B. O’Hara, P. Solymos, M. H. H. Stevens, E. Szöcs, H. H. Wagner, M. Barbour, M. Bedward, B. Bolker, D. Borcard, G. Carvalho, M. Chirico, M. De Caceres, S. Durand, H. B. A. Evangelista, R. FitzJohn, M. Friendly, B. Furneaux, G. Hannigan, M. O. Hill, L. Lahti, D. McGlinn, M.-H. Ouellette, E. Ribeiro Cunha, T. Smith, A. C. Stier, C. J. F. Ter Braak, and J. Weedon. 2022. vegan: Community ecology package R package version 2.6-2.

Penin, L., F. Michonneau, A. H. Baird, S. R. Connolly, M. S. Pratchett, M. Kayal, and M. Adjeroud. 2010. “Early post-settlement mortality and the structure of coral assemblages.” Marine Ecology Progress Series 408: 55–64. https://doi.org/10.3354/meps08554.

Privitera-Johnson, K., E. A. Lenz, and P. J. Edmunds. 2015. “Density-associated recruitment in octocoral communities in St. John, US Virgin Islands.” Journal of Experimental Marine Biology and Ecology 473: 103–109. https://doi.org/10.1016/j.jembe.2015.08.006.

R Core Team. 2022. R: A language and environment for statistical computing 4.2.1. R Foundation for Statistical Computing, Vienna, AT.

Rasher, D. B., E. P. Stout, S. Engel, J. Kubanek, and M. E. Hay. 2011. “Macroalgal terpenes function as allelopathic agents against reef corals.” Proceedings of the National Academy of Sciences 108 (43): 17726. https://doi.org/10.1073/pnas.1108628108.

River, G. F., and P. J. Edmunds. 2001. “Mechanisms of interaction between macroalgae and scleractinians on a coral reef in Jamaica.” Journal of Experimental Marine Biology and Ecology 261 (2): 159–172. https://doi.org/10.1016/S0022-0981(01)00266-0.

Roff, G., and P. J. Mumby. 2012. “Global disparity in the resilience of coral reefs.” Trends in Ecology & Evolution 27 (7): 404–413. https://doi.org/10.1016/j.tree.2012.04.007.

Ruzicka, R. R., M. A. Colella, J. W. Porter, J. M. Morrison, J. A. Kidney, V. I. P. Brinkhuis, K. S. Lunz, K. A. Macaulay, L. A. Bartlett, M. K. Meyers, and J. Colee. 2013. “Temporal changes in benthic assemblages on Florida Keys reefs 11 years after the 1997/1998 El Niño.” Marine Ecology Progress Series 489: 125–141. https://doi.org/10.3354/meps10427.

Sammarco, P. W. 1980. “Diadema and its relationship to coral spat mortality: Grazing, competition, and biological disturbance.” Journal of Experimental Marine Biology and Ecology 45 (2): 245–272. https://doi.org/10.1016/0022-0981(80)90061-1.

Sánchez, J. A., M. Gómez-Corrales, L. Gutierrez-Cala, D. C. Vergara, P. Roa, F. L. González-Zapata, M. Gnecco, N. Puerto, L. Neira, and A. Sarmiento. 2019. “Steady decline of corals and other benthic organisms in the SeaFlower Biosphere Reserve (Southwestern Caribbean).” Frontiers in Marine Science 6: 1–13. https://doi.org/10.3389/fmars.2019.00073.

Sebens, K. P. 1983a. “The larval and juvenile ecology of the temperate octocoral Alcyonium siderium Verrill. I. Substratum selection by benthic larvae.” Journal of Experimental Marine Biology and Ecology 71 (1): 73–89. https://doi.org/10.1016/0022-0981(83)90105-3.

Sebens, K. P. 1983b. “The larval and juvenile ecology of the temperate octocoral Alcyonium siderium Verrill. II. Fecundity, survival, and juvenile growth.” Journal of Experimental Marine Biology and Ecology 72: 263–285. https://doi.org/10.1016/0022-0981(83)90111-9.

Sebens, K. P. 1986. “Spatial relationships among encrusting marine organisms in the New England subtidal zone.” Ecological Monographs 56 (1): 73–96. https://doi.org/10.2307/2937271.

Slattery, M., G. A. Hines, J. Starmer, and V. J. Paul. 1999. “Chemical signals in gametogenesis, spawning, and larval settlement and defense of the soft coral Sinularia polydactyla.” Coral Reefs 18 (1): 75–84. https://doi.org/10.1007/s003380050158.

Stockton, L., and P. J. Edmunds. 2021. “Spatially aggressive peyssonnelid algal crusts (PAC) constrain coral recruitment to Diadema grazing halos on a shallow Caribbean reef.” Journal of Experimental Marine Biology and Ecology 541: 151569. https://doi.org/10.1016/j.jembe.2021.151569.

Therneau, T. M. 2022. A package for survival analysis in R R package version 3.3-1.

Tonra, K. J., C. D. Wells, and H. R. Lasker. 2021. “Spawning, embryogenesis, settlement, and post-settlement development of the gorgonian Plexaura homomalla.” Invertebrate Biology: e12319. https://doi.org/10.1111/ivb.12319.

Tsounis, G., and P. J. Edmunds. 2017. “Three decades of coral reef community dynamics in St. John, USVI: A contrast of scleractinians and octocorals.” Ecosphere 8 (1): e01646. https://doi.org/10.1002/ecs2.1646.

Tsounis, G., M. A. Steele, and P. J. Edmunds. 2020. “Elevated feeding rates of fishes within octocoral canopies on Caribbean reefs.” Coral Reefs 39: 1299–1311. https://doi.org/10.1007/s00338-020-01963-1.

Venables, W. N., and B. D. Ripley. 2002. Modern applied statistics with S. Fourth edition ed. New York, NY, USA: Springer.

Vreeland, H. V., and H. R. Lasker. 1989. “Selective feeding of the polychaete Hermodice carunculata Pallas on Caribbean gorgonians.” Journal of Experimental Marine Biology and Ecology 129 (3): 265–277. https://doi.org/10.1016/0022-0981(89)90108-1.

Wells, C. D., Á. Martínez-Quintana, K. J. Tonra, and H. R. Lasker. 2021. “Algal turf negatively affects recruitment of a Caribbean octocoral.” Coral Reefs 40: 1045–1053. https://doi.org/10.1007/s00338-021-02103-z.

Williams, S. M. 2022. “The reduction of harmful algae on Caribbean coral reefs through the reintroduction of a keystone herbivore, the long-spined sea urchin Diadema antillarum.” Restoration Ecology 30 (1): e13475. https://doi.org/10.1111/rec.13475.

Wolf, A. T., and M. M. Nugues. 2013. “Predation on coral settlers by the corallivorous fireworm Hermodice carunculata.” Coral Reefs 32 (1): 227–231. https://doi.org/10.1007/s00338-012-0969-x.

Wood, S. N. 2021. Mixed GAM computation vehicle with automatic smoothness estimation R package version 1.8-38.

Yoshioka, P. M. 1994. “Size-specific life history pattern of a shallow-water gorgonian.” Journal of Experimental Marine Biology and Ecology 184 (1): 111–122. https://doi.org/10.1016/0022-0981(94)90169-4.

Yoshioka, P. M. 1996. “Variable recruitment and its effects on the population and community structure of shallow-water gorgonians.” Bulletin of Marine Science 59 (2): 433–443.

Yoshioka, P. M., and B. B. Yoshioka. 1989. “Effects of wave energy, topographic relief and sediment transport on the distribution of shallow-water gorgonians of Puerto Rico.” Coral Reefs 8 (3): 145–152. https://doi.org/10.1007/BF00338270.

