## Supplementary Figures 1-4 and Table 1-2 for "Grazers and predators mediate the post-settlement bottleneck in Caribbean octocoral forests"

**Supplementary Materials**

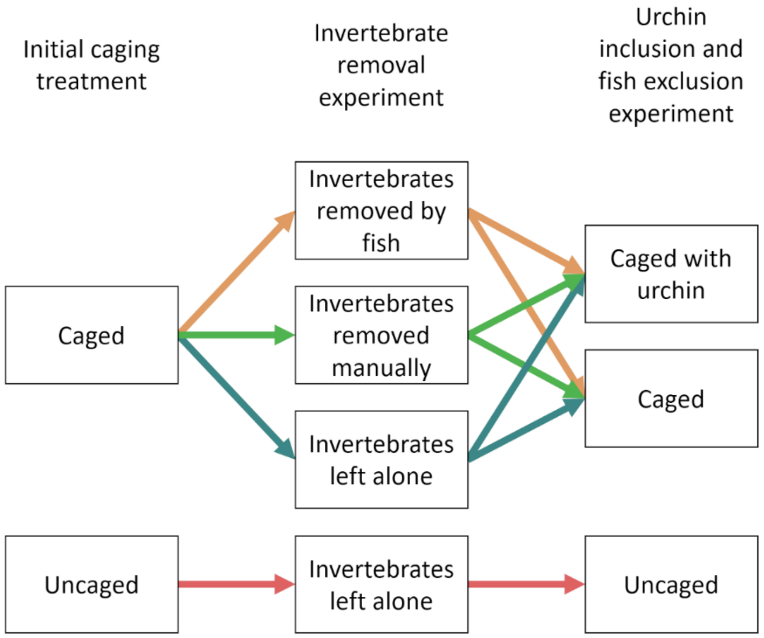


Supplemental Figure 1. Schematic for chronology of the experimental design.

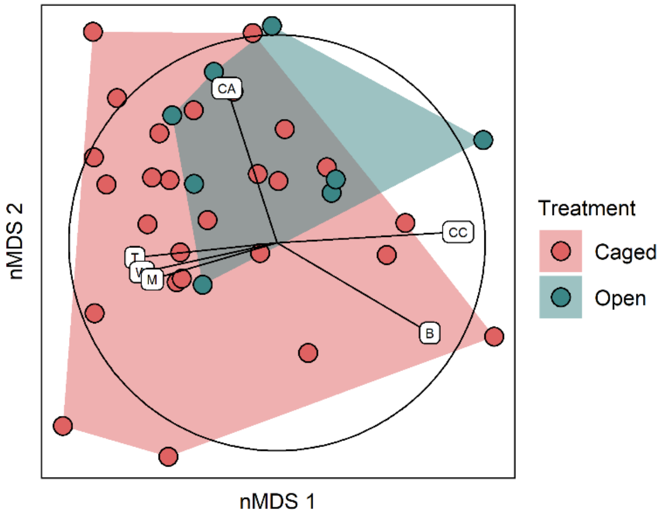


Supplemental Figure 2. Non-metric multidimensional scaling plot of the tile cover communities on the bottom of the tiles with vectors displaying Spearman coefficients between cover types and corresponding nMDS scores. The circle represents a Spearman correlation of 1.0 and vectors are a fraction of that unit circle. Only cover types with a correlation of over 0.6 with the scores of the first and second dimensions are displayed. **B**: bare tile; **CA**: colonial ascidians; **CC**: crustose coralline algae; **M**: macroalgae; **T**: turf algae; **W**: calcified worm tubes.

Supplemental Table 1. Average number of invertebrates removed from manual removal tiles. Bolded taxa are phyla and indented taxa are subgroups within their phyla.

| Taxa | No. of individuals removed |
|---|---|
| **Annelida** | **106** |
| Errantia | 35 |
| Sedentaria | 71 |
| **Arthropoda** | **72** |
| Amphipoda | 53 |
| Copepoda | 6 |
| Cephalocarida | 0 |
| Decapoda | 4 |
| Isopoda | 1 |
| Mysida | 4 |
| Ostracoda | 5 |
| **Mollusca** | **17** |
| Bivalvia | 4 |
| Gastropoda | 13 |
| **Nematoda** | **10** |
| *Total* | *195* |

Supplemental Table 2. Percent difference of invertebrate counts after the mesofauna removal experiment compared to invertebrates removed in the manual removal treatment before the experiment. Bolded taxa are phyla and indented taxa are subgroups within their phyla.

| Taxa | No removal | Fish removal | Manual removal | Reef removal |
|---|---|---|---|---|
| **Annelida** | **-33 %** | **-53 %** | **-75 %** | **-84 %** |
| Errantia | -11 % | -20 % | -29 % | -63 % |
| Sedentaria | -44 % | -69 % | -97 % | -94 % |
| **Arthropoda** | **-83 %** | **+2 %** | **-88 %** | **-53 %** |
| Amphipoda | -87 % | -45 % | -92 % | -70 % |
| Copepoda | -83 % | +567 % | -50 % | -33 % |
| Cephalocarida | -100 % | +500 % | -100 % | -100 % |
| Decapoda | -75 % | -75 % | -100 % | -100 % |
| Isopoda | -100 % | -100 % | -100 % | +1250 % |
| Mysida | -50 % | -75 % | -100 % | 0 % |
| Ostracoda | -79 % | -79 % | -57 % | -79 % |
| **Mollusca** | **-88 %** | **-34 %** | **-88 %** | **-100 %** |
| Bivalvia | -100 % | -100 % | -100 % | -100 % |
| Gastropoda | -85 % | -15 % | -85 % | -100 % |
| **Nematoda** | **-90 %** | **+86 %** | **+117 %** | **-79 %** |
| *Total* | *-56 %* | *-31 %* | *-81 %* | *-74 %* |

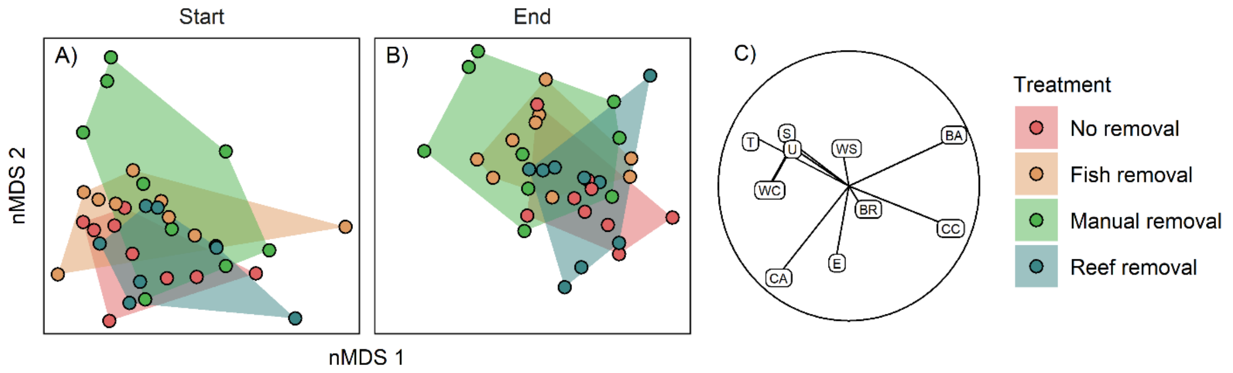


Supplemental Figure 3. Non-metric multidimensional scaling plot of the tile cover communities on the (A-B) bottom of the tiles alongside (C) Spearman coefficients between cover types and corresponding nMDS at the (A) start and (B) end of the experiment. The circle represents a Spearman correlation of 1.0 and vectors are a fraction of that unit circle. Only cover types with a correlation of over 0.2 with the scores of the first and second dimensions are displayed. **BA**: bare tile; **BR**: bryozoans; **CA**: colonial ascidians; **CC**: crustose coralline algae; **E**: encrusting algae; **S**: sponge; **T**: turf algae; **U**: upright algae; **WC**: calcified worm tubes; **WS**: proteinaceous worm tubes.

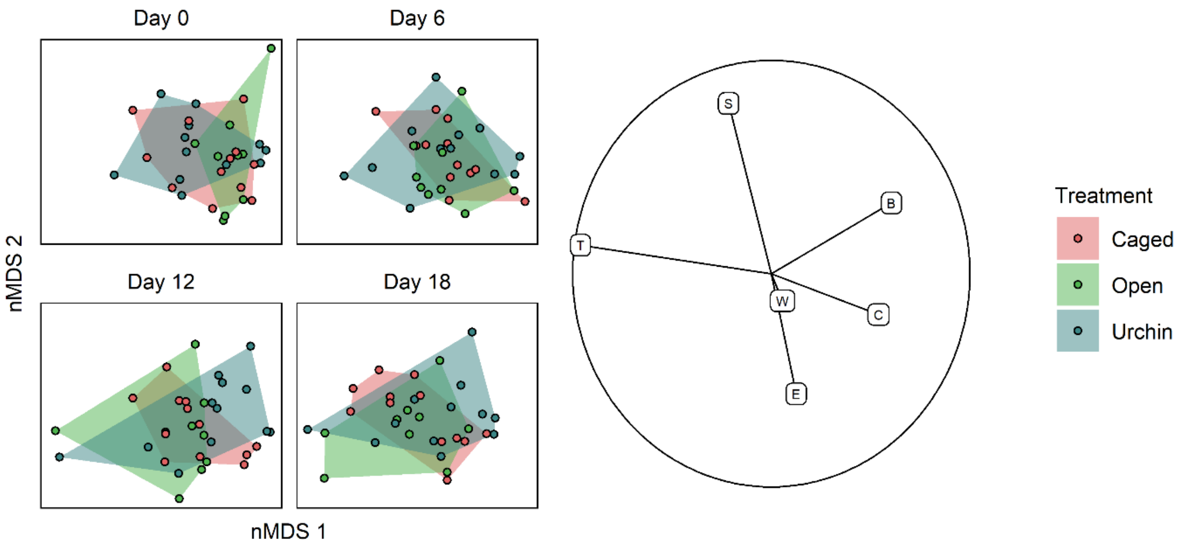


Supplemental Figure 4. Non-metric multidimensional scaling plot of the tile cover communities on the bottom of the tiles alongside Spearman coefficients between cover types and corresponding nMDS. The circle represents a Spearman correlation of 1.0 and vectors are a fraction of that unit circle. **C**: crustose coralline algae; **T**: turf algae; **B**: bare tile; **S**: sponge; **E**: encrusting algae; **W**: calcified worm tubes.
